# LSD Relaxes Structural Constraints on Brain Dynamics: A Connectome Harmonic MEG Study

**DOI:** 10.64898/2026.03.02.709138

**Authors:** Venkatesh Subramani, Annalisa Pascarella, Jérémy Brunel, Yann Harel, Suresh Muthukumaraswamy, Robin Carhart-Harris, Karim Jerbi, Giulia Lioi, Nicolas Farrugia

**Affiliations:** Cognitive and Computational Neuroscience Laboratory (CoCo Lab), Psychology Department, Université de Montréal, Québec, Canada; IMT Atlantique, Lab-STICC, UMR CNRS 6285, F-29238, Brest, France; Institute of Applied Mathematics ”M.Picone” -CNR, Italy; Centre de recherche Hôpital Maisonneuve-Rosemont (HMR), Montreal, QC, Canada; CRIUGM (Centre de Recherche de l’Institut Universitaire de Gériatrie de Montréal), Montréal, QC, Canada; University of Auckland, Auckland, New Zealand; University of California, San Francisco, San Francisco, United States; Mila (Quebec AI Institute), Montréal, Québec, Canada; UNIQUE Center (Quebec Neuro-AI Research Center), Montréal, Québec, Canada

**Keywords:** Consciousness, Psychedelics, Lysergic acid diethylamide (LSD), Magnetoencephalography (MEG), Brain structure-function relationship, Graph analysis, Diffusion-weighted imaging

## Abstract

Psychedelics profoundly alter conscious experience, yet how they reshape the relationship between brain anatomy and electrophysiological dynamics remains unclear. Here we use source-localized magnetoencephalography mapped onto connectome harmonics to quantify structure–function coupling in humans under lysergic acid diethylamide (LSD) and placebo. LSD induces a robust decoupling of low-frequency (theta, alpha and beta) activity from anatomical constraints, indicating a global loosening of structure-aligned large-scale dynamics. High-frequency gamma activity shows selective reorganization rather than uniform disruption. Decoupling within core default-mode network regions predicts ego dissolution intensity across individuals, linking frequency-selective DMN reorganization to subjective loss of self. Functional decoding further reveals system-specific rebalancing: visual and attentional systems preferentially decouple while auditory networks exhibit strengthened coupling. Together, these findings provide electrophysiological evidence that psychedelic states emerge from a frequency-dependent relaxation of structural constraints on brain activity and identify default-mode reorganization as a neural correlate of ego dissolution. These results offer a mechanistic framework for understanding how LSD may exert therapeutic effects by transiently relaxing rigid structural constraints and enhancing dynamical flexibility within networks involved in self-related processing.

## 1 Introduction

Just as a violin generates a rich repertoire of melodies through physical constraints imposed by its strings and body, the brain’s anatomical substrate enables a wide range of brain states and, in turn, conscious experience [1]. Psychedelics provide a powerful lever for perturbing brain dynamics, reliably inducing altered states of consciousness characterized by perceptual distortions, enhanced imagery, affective changes, and ego dissolution [2, 3]. The relationship between brain structure and function has increasingly been viewed as a fundamental axis along which levels and contents of consciousness can be characterized [4]. This perspective motivates the present study, which probes how LSD modulates the constraints imposed by anatomical structure on electrophysiological brain dynamics.

Traditionally, structure–function coupling (SFC) has been studied by relating structural connectomes (SC) to functional connectivity (FC), revealing that FC is partly predictable from underlying anatomy [5]. More recent work has formalized this relationship using predictive models, including deep learning approaches that infer individual FC from SC [6]. However, SC–FC correspondence does not directly capture how functional activity itself *aligns* with anatomy, in the sense of how neural signals are spatially organized and constrained by the structural connectome. Addressing this question is enabled by connectome harmonics (CH), which extend Fourier analysis to signals defined on the brain’s network architecture [7].

In the temporal domain, signals are decomposed into harmonic modes (sinusoidal waves) ordered by temporal frequency. Analogously, CH re-expresses functional brain activity in terms of spatial harmonics of the structural connectome, ordered by spatial frequency [7]. Similar to how any temporal activity can be represented using varying frequencies of sinusoidal waves, CH allows any cortical activity pattern to be captured as a weighted combination of connectome-derived spatial modes. Spatial frequency of the modes (eigenmodes) holds a particular relevance. The spatially smooth eigenmodes (low-order) express the functional activity that propagates along the underlying structural architecture, interpreted as reflecting strong structure–function *coupling*. In contrast, spatially localized eigenmodes (high-order) characterize functional activity that is less constrained by anatomical connectivity, interpreted as reflecting *decoupling* from structure.

Using connectome harmonics, prior work has characterized the organization of SFC during normal wakefulness [8, 9, 10, 11, 12, 13, 14, 15]. These studies converge in showing strong structure–function alignment in unimodal cortices, while transmodal regions exhibit more heterogeneous coupling–decoupling profiles that differ across electrophysiological and hemodynamic activity. These modality-specific differences highlight that structure–function relationships are intrinsically shaped on the temporal scale and biophysical origin of the measured signal.

Psychedelics are proposed to increase the diversity and flexibility of neural states [2, 3]. fMRI-based connectome harmonic analyses suggest that psychedelics redistribute cortical activity pattern from spatially smooth pattern to distributed patterns [16, 17, 18]. Because the higher-order distributed harmonics capture the brain activity that reflects relaxation of structural constraints, this shift (i.e. higher weights) has been interpreted as reflecting increased entropy, and expanded functional repertoire. However, such inferences rely on hemodynamic signals, which may dissociate from underlying neural activity, particularly under psychedelics [19, 20]. As a result, psychedelic-induced changes observed in fMRI may reflect not only neural reorganization but also vascular modulation. Moreover, cortical dynamics are inherently frequency-specific: low-frequency rhythms emerge from large-scale coordination of distributed neural populations, whereas high-frequency activity reflects more local circuit processes [21, 22]. How anatomical structure constrains neural activity across frequencies therefore cannot be resolved using hemodynamic measures alone.

Here, for the first time, we examine psychedelic-induced reorganization of structure–function coupling using source-localized MEG, which provides a direct, millisecond-resolved measure of neural activity. Leveraging connectome harmonics, we map source-localized MEG activity onto the structural connectome to quantify the extent to which neural dynamics align with anatomical pathways under LSD relative to placebo (PLA). We hypothesize that LSD induces a frequency-dependent reconfiguration of structure–function coupling, selectively relaxing anatomical constraints on slow, integrative dynamics while reshaping fast, locally expressed activity. This approach offers a spatial description of how psychedelics reorganize brain function at the level of neural dynamics, rather than relying on indirect metabolic proxies.

## 2 Materials and Methods

### 2.1 Data

We analyzed MEG data collected by Carhart-Harris and co. [23]. The original study followed a single-blind, within-subject, placebo-controlled setup in which 17 participants received intravenous LSD (75 *µ*g) or placebo in counterbalanced order across two sessions separated by 14 days. MEG was acquired 4 h post-administration during eyes-closed rest with (Music) or without music (NoMusic; 7 min per condition). Further details are reported elsewhere [23, 24].

### 2.2 Preprocessing

MEG preprocessing followed the pipeline of the original dataset [23], augmented with automated artefact detection using Autoreject [25] (Supp. Fig. S1). Continuous recordings were epoched into non-overlapping 2-s segments and cleaned using a combination of Autoreject and independent component analysis to attenuate physiological and non-physiological artefacts (See Supplementary Material for details).

### 2.3 Source localization

Source localization was performed using individual T1-weighted MRI scans processed with FreeSurfer. Forward models were computed using boundary element method, and source estimates were obtained using dynamic statistical parametric mapping (dSPM). Individual source estimates were morphed to a common template (fsaverage) and parcellated into 360 cortical regions using the HCP-MMP atlas [26]. Source-level power spectral density was estimated using a multitaper approach and served as the basis for the spectral analysis. Structure–function coupling was quantified for the bandlimited activity (butterworth filter, order of 4) for theta (4-8 Hz) through high-gamma (90-120 Hz) range. Full source reconstruction parameters are reported in the Supplementary Methods.

### 2.4 Structure-Function Coupling

#### 2.4.1 Structural connectome

Because diffusion-weighted imaging was not available for the MEG dataset, we derived the structural connectome from high-quality diffusion MRI data provided by the Human Connectome Project (HCP S1200 release; *N* = 1,063) [27]. Individual connectivity matrices were reconstructed and averaged to obtain a consensus connectome. Edge weights were defined as streamline density between pairs of cortical regions parcellated according to the HCP-MMP atlas, matching the atlas used for source-localized MEG activity. To obtain a group-level connectome suitable for our analyses, individual connectivity matrices were averaged across participants to form a consensus structural connectome (*N* = 1,063) (Fig. 1A). Full tractography and reconstruction details are provided in the Supplementary Methods.

**Figure 1:**
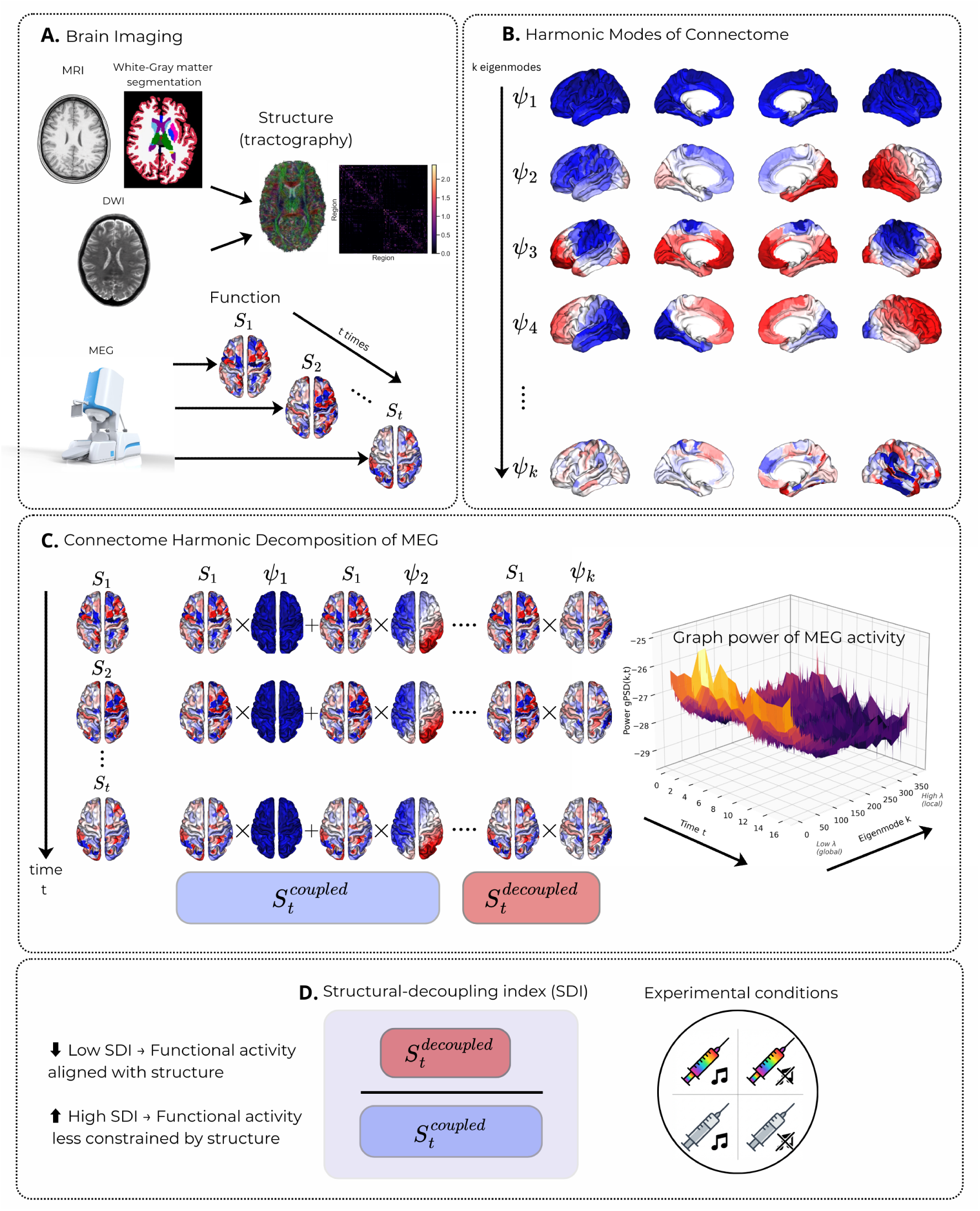
Overview of Methodology. **A** Anatomical MRI and diffusion-weighted imaging are used to segment white and gray matter and to reconstruct an anatomically constrained tractography. Streamline densities between cortical regions defined by the HCP-MMP atlas are used to build a group-level structural connectome. Source-localized MEG provides time-resolved cortical activity *S_t_*. **B** Its harmonic modes *ψ_k_*are obtained via eigendecomposition of the graph Laplacian (See Supplementary). Low-order eigenmodes capture spatially smooth, global patterns constrained by anatomy, whereas high-order modes capture increasingly localized patterns weakly constrained by structure. **C** Cortical MEG activity is decomposed at each time point as a weighted linear combination of connectome harmonic modes, yielding a representation of functional activity in the graph spectral domain. The resulting graph power spectral density (gPSD) quantifies the contribution of each harmonic mode over time. Graph-domain filtering (median split of the gPSD) separates structure-coupled and structure-decoupled components of the signal. **D** The relative balance between decoupled and coupled components defines the Structural-Decoupling Index (SDI), which is computed across experimental conditions (LSD vs placebo; music vs no-music) to quantify changes in structure–function coupling (See Supplementary). MEG system image credits to CTF MEG Neuro Innovations, Inc.

#### 2.4.2 ConnectomeHarmonic Decomposition and Structural Decoupling Index

To characterize how electrophysiological activity aligns with anatomical structure, we combined source-localized MEG with connectome harmonic analysis (Fig. 1). A consensus structural connectome derived from diffusion MRI defines an intrinsic set of spatial harmonics spanning smooth, structure-aligned patterns to increasingly localized patterns that deviate from large-scale anatomical constraints (Fig. 1B). Source-level MEG activity was projected onto this harmonic basis, allowing cortical dynamics to be expressed as a weighted combination of structure-coupled and structure-decoupled components (Fig. 1C). We quantified the relative dominance of these components using the Structural Decoupling Index (SDI [8]), which captures the ratio between activity aligned with the structural connectome and activity expressed in higher-order, spatially localized modes (Fig 1D). Following [8], binary log is subsequently applied, resulting in the negative and positive tail corresponding to coupling and decoupling respectively. This framework provides an anatomically grounded measure of structure–function coupling that is directly applicable to fast electrophysiological signals. Full mathematical definitions and implementation details are provided in the Supplementary Methods.

Consistent with prior studies under normal wakeful state [9, 14], we computed SDI on broadband cortical signals, without separating periodic oscillatory activity from the aperiodic (1/*f* -like) component. This allows for a direct comparison of electrophysiological SFC across normal wakeful state and psychedelics-induced altered states.

### 2.5 Functional correlates of spatial maps

To interpret LSD-induced changes in structure–function coupling at the systems level, we investigated the functional correlates of SDI contrast maps using NiMARE [28], following established approaches in prior work [29, 8, 14]. Unthresholded SDI maps were compared against the Neurosynth meta-analytic database to identify large-scale cognitive and affective domains associated with regions exhibiting the strongest coupling and decoupling effects under LSD. This approach enables a systems-level interpretation of structure–function reorganization by relating spatial patterns of SDI change to distributed functional domains (e.g., perception, attention, emotion, language), rather than to individual tasks or regions. Full decoding procedures and statistical thresholds are detailed in the Supplementary Methods.

### 2.6 Phenomenology regression

We examined the correspondence between changes in SDI and phenomenological scores using Spearman’s correlation coefficient *ρ*. To this end, the regression analysis were performed within *a priori* regions or functional networks, selected based on previous studies. Multiple comparisons were controlled using Bonferroni correction across number of spatial tests. The 95% CI was established using bias-corrected bootstrapping [30] technique (N = 10,000) implemented in scipy.stats.bootstrap.

### 2.7 Statisticalframework

We employed cluster-based permutation testing (mne.stats.permutation_cluster_test) to assess the statistical significance of changes in graph power and SDI. For gPSD, clusters were defined along the one-dimensional eigenmode axis, reflecting adjacency in graph-frequency space. For SDI, clusters were defined based on spatial adjacency among HCP-MMP parcels. Cluster-level significance was evaluated at *α* = 0.05.

The *Main Effect* of Drug and Music assessed the impact of each factor independently of the other. The *Interaction effect (Drug × Music)* tested whether the effect of LSD relative to placebo differed as a function of music presence.

## 3 Results

Using connectome harmonics as the basis of brain activity, we characterize how LSD reorganizes the structural constraints on brain activity, mapping these shifts to large-scale functional systems and subjective phenomenology.

### 3.1 Absolutegraph power changes mirror spectral power changes

LSD-induced modulations (i.e. Main Effect Drug) in the spatial organization of cortical activity were revealed by mapping the source-localized cortical activity onto the structural connectome (Fig. 2). The resulting graph power (gPSD) describes the distribution of the weights of each mode along the spatial scales ranging from smooth, large-scale patterns (low eigenvalues) to more spatially localized patterns (high eigenvalues).

**Figure 2:**
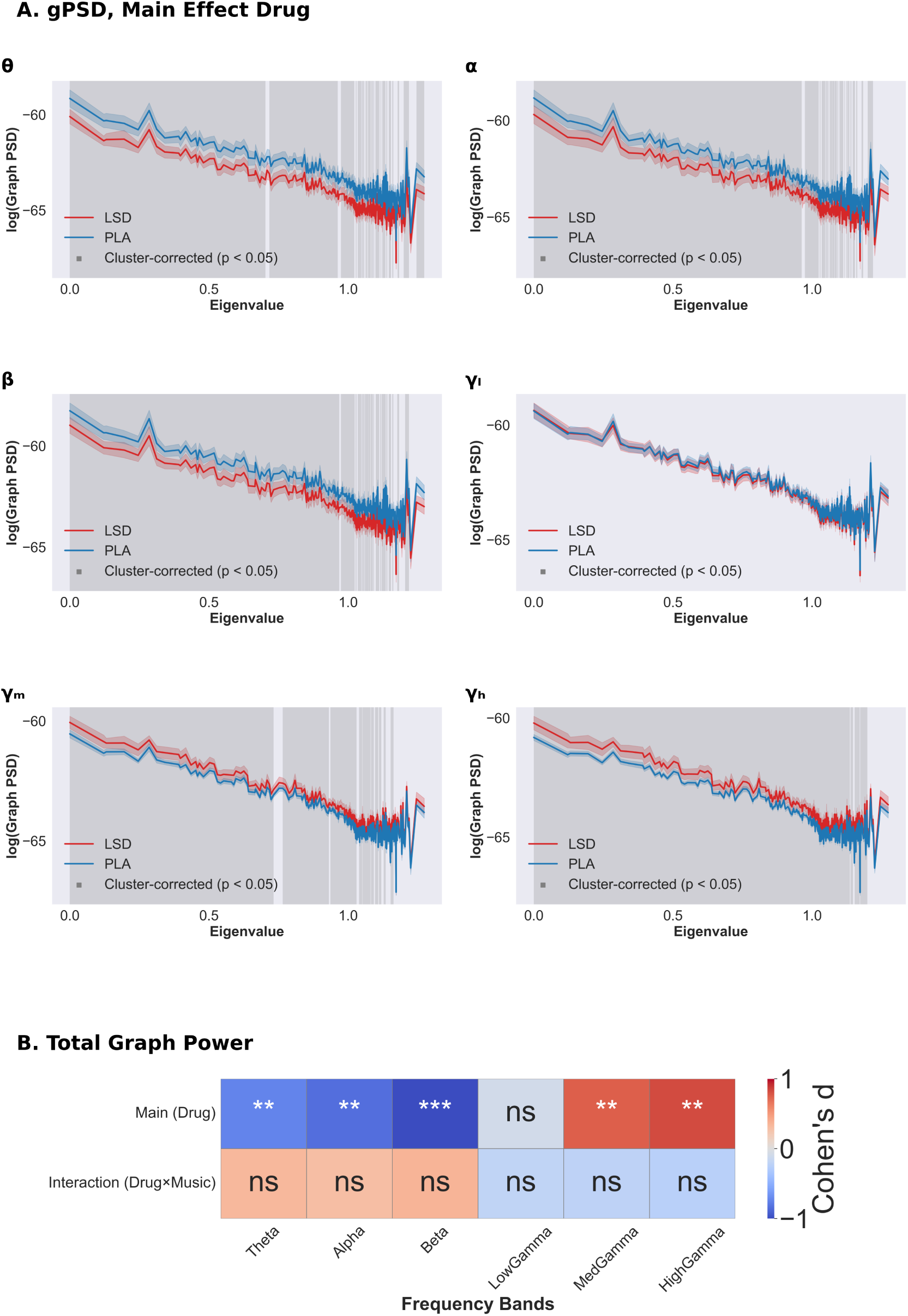
Absolute power reductions and increases in the graph spectral domain. **A** LSD-induced changes in the graph Power Spectral Density (gPSD) across canonical frequency ranges: *θ* (4 -8 Hz), *α* (8 -13 Hz), *β* (15 -30 Hz), low-*γ* (30 -60 Hz), mid-*γ* (60 -90 Hz) and high-*γ* (90 -120 Hz). The gPSD characterizes the graph spectral power of the cortical activity on the structural graph domain as a function of spatial frequencies. Red line indicates graph spectral power during LSD condition. Significance differences (Cluster-tests, *p <* 0.05, N = 10,000 permutations) are highlighted in grey shade. **B** Total Graph Power changes for the effect of drug and the effect of interaction between drug and music over frequency bands. Significance level established using paired t-tests, and the cells are colorcoded by the effect size (Cohen’s d). * / ** / *** correspond to *p <* 0.05 / *p <* 0.01 / *p <* 0.001

Significant changes (*p <* 0.05, cluster-corrected, *N* = 10,000 permutations) in graph power were observed across all frequency bands except low-gamma (Fig. 2A). Significantly weakened graph power was observed in theta through beta range across a broad range of eigenmodes (Fig 1B for the spatial topography). In contrast, mid- and high-gamma bands showed increased graph power, with effects concentrated in low- to mid-range eigenmodes, corresponding to large-scale and intermediate spatial patterns along the structural connectome.

To summarize these effects, we computed total graph power by summing gPSD across all eigenmodes. Total graph power was examined for the main effect of drug and the drug × music interaction using paired t-tests, with effect sizes visualized using Cohen’s *d* (Fig. 2B). The main drug effect condition revealed a broadband modulation of total graph power, with the strongest effect observed in the beta band. No frequency band exhibited a statistically significant drug × music interaction. Nevertheless, effect-size estimates showed a consistent reversal in the sign of Cohen’s *d* across several frequency bands, suggesting a systematic attenuation of LSD-induced power changes in the presence of music, although this trend did not reach statistical significance.

### 3.2 LSD decouples theta, alpha and beta activity from the structure

With the global reduction in the graph power observed, characterizing the relative proportion of the graph power between localized eigenmodes and smooth eigenmodes revealed the SFC organization. Group-level maps (*p <* 0.05, cluster-corrected, *N* = 10, 000 permutations) for the main effect of drug are shown in Fig. 3. Positive values (red) indicate LSD-induced decoupling of functional activity from the underlying structural architecture relative to PLA, whereas negative values (blue) indicate stronger structure–function coupling under LSD.

**Figure 3:**
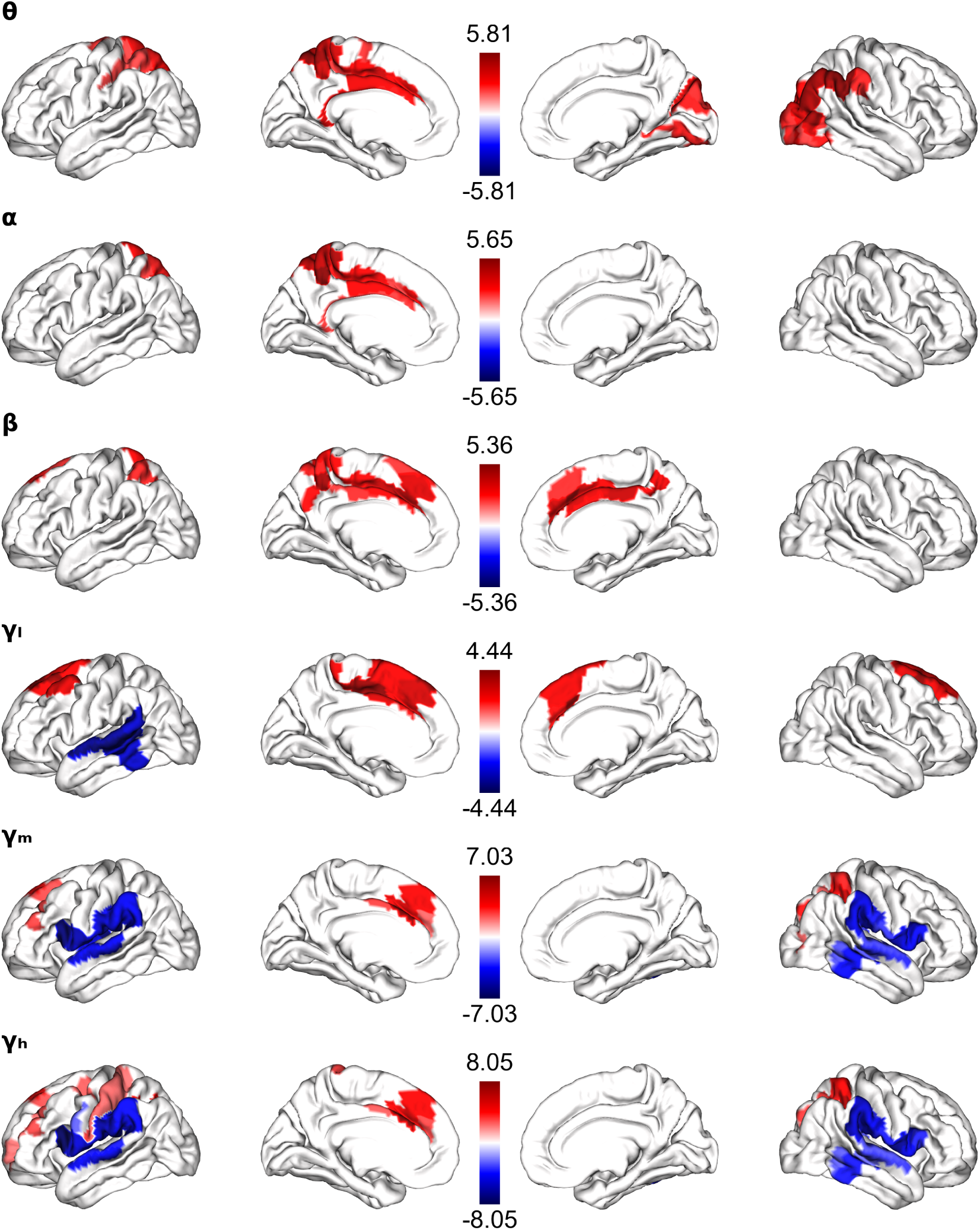
LSD-induced changes in structure-function relationship. Structure-function coupling and decoupling quantified for the temporal signals across canonical frequency ranges. Significance level was assessed using cluster-tests (LSD vs PLA, *p <* 0.05, N = 10,000 permutations) and the red (blue) tail in these t-maps corresponds to LSD-induced decoupling (coupling) of the functional activity from the structure.

In the low-frequency range (theta-beta) LSD induced decoupling (relative to PLA), with effects that were spatially focal yet consistent across bands. In contrast, higher frequencies exhibit a more heterogeneous pattern, with both significant coupling and decoupling observed depending on cortical region. Quantifying the spatial similarity across frequency bands using Spearman’s *ρ* revealed a strong correspondence among theta, alpha, beta maps (*ρ* > 0.82, *p <* 10^−15^; Supplementary figure S3), shedding light on a coherent low-frequency mode of LSD-induced decoupling.

In the high-frequency range, alongside regions of decoupling, we observed increases in structure–function coupling, most prominently localized to temporal cortical areas. Spatial similarity analysis revealed mid- and high-gamma were near identical (*ρ* = 0.98, *p <* 10^−15^), highlighting a distinct and consistent high-frequency mode of structural reorganization. Low-gamma occupied an intermediate position, showing moderate similarity to both low- and high-frequency bands, and thus represents a transitional regime separating these two clusters.

### 3.3 Functionalcorrelates reveals system-specific perturbations of LSD

To assess the functional relevance of LSD-induced reorganization in SFC, we mapped the SFC changes onto large-scale cognitive and affective domains using NiMARE (Fig. 4). This analysis focused on the main effect of drug (Fig. 4). In an exploratory analysis, we additionally examined the drug × music interaction (Supp. Fig. S7A), and the main effect of music (Supp. Fig. S7B), enabling assessment of how distinct experimental factors modulate SFC organization and its functional relevance.

**Figure 4:**
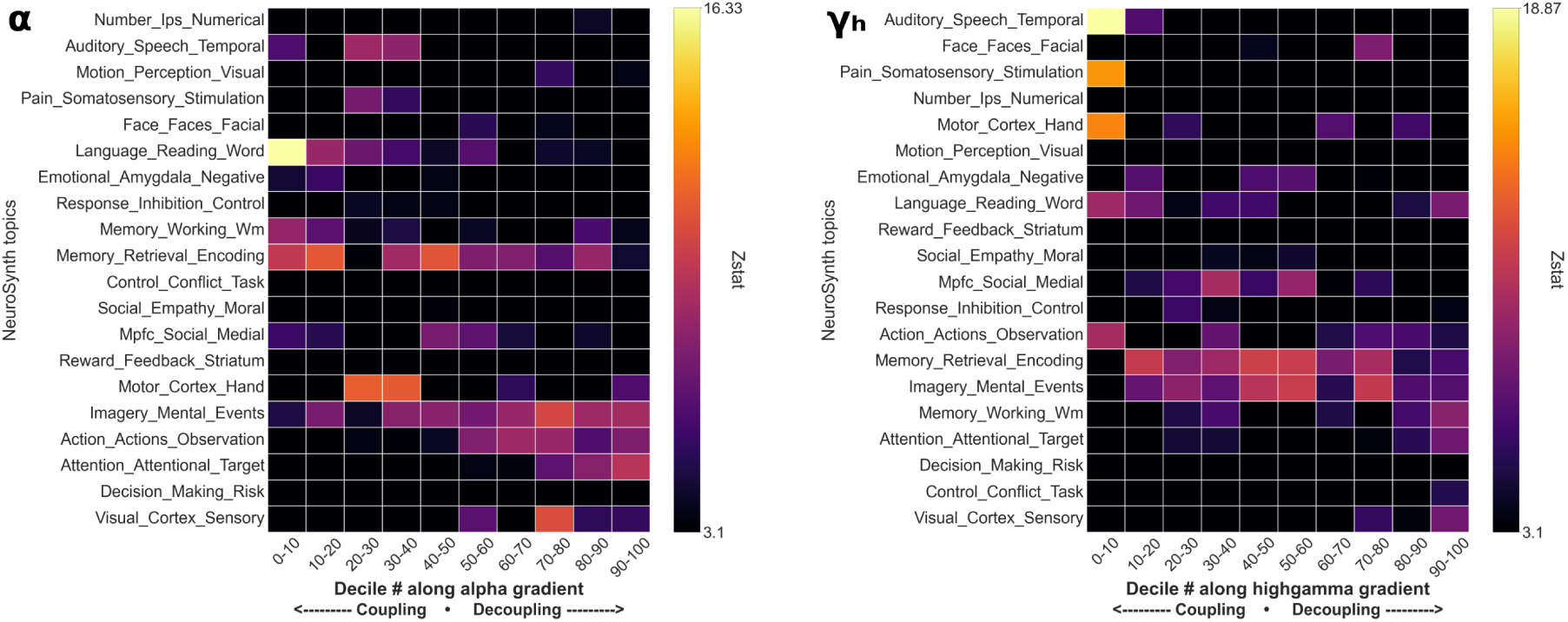
Meta-analytic functional correlates of LSD-induced SFC reorganization. SDI contrast maps for the main effect of drug in the *α* (left) and high-*γ* (right) bands. T-maps were binned into deciles, binarized, and submitted to NiMARE for meta-analytic functional association. For each decile, spatial correlations between the SDI-derived masks and meta-analytic topic maps were computed. Correlation values were *z*-scored and thresholded at *p <* 0.0001.

Functional domains associated with LSD-induced SDI changes are summarized in Fig. 4. The horizontal axis represents a gradient from stronger structure–function coupling (left) to stronger decoupling (right) under LSD relative to PLA. Across alpha and high-gamma bands, the spatial topography of effects was consistent in some functional systems but diverged in others. Notably, the auditory system exhibited stronger coupling most prominently in the high-gamma band. Furthermore, LSD-induced strengthening of coupling within auditory, language, and emotion domains was more pronounced in alpha band than in high-gamma band. In contrast, across both frequency bands, domains associated with visual processing, executive control, action, and attention are preferentially positioned toward the decoupling end of the gradient.

Exploratory analyses of the drug × music interaction revealed a marked reversal in several functional systems (Supplementary Fig. S7A). Functional domains that exhibited LSD-induced structure–function coupling in the main drug effect shifted toward decoupling in the presence of music, with the strongest reversals observed in auditory and emotion-related systems. Conversely, functional systems that exhibited prominent LSD-induced decoupling, such as visual and attention-related domains, shifted toward stronger coupling when LSD was administered in conjunction with music. Finally, assessing the functional correlates of main effect of music independent of drug state (Music-NoMusic SDI contrasts; Supplementary Fig. S7B) revealed that music was associated with structure–function decoupling in auditory, emotional, and visual cortices, indicating reduced alignment with underlying structural constraints during music listening irrespective of pharmacological condition.

### 3.4 Changes in structure-function coupling relate to phenomenology

The relationship between reorganization in SFC and subjective experience was assessed by computing Spearman’s correlation coefficient (*ρ*) between SDI values and phenomenological dimensions. Four phenomenological dimensions were examined : Complex Imagery, Ego Dissolution, Emotional Arousal and Positive Mood. These associations were evaluated separately for the main effect of drug, and the interaction between drug and music. Figure 5 presents the significant correlation between changes in SFC and phenomenology (*p <* 0.05, Bonferroni-corrected for number of spatial tests). Regression analyses were performed within a subset of *a priori* regions selected based on prior literature. Heatmaps display the correlation coefficients, while the spatial maps above panel label indicate regions included in each analysis.

**Figure 5:**
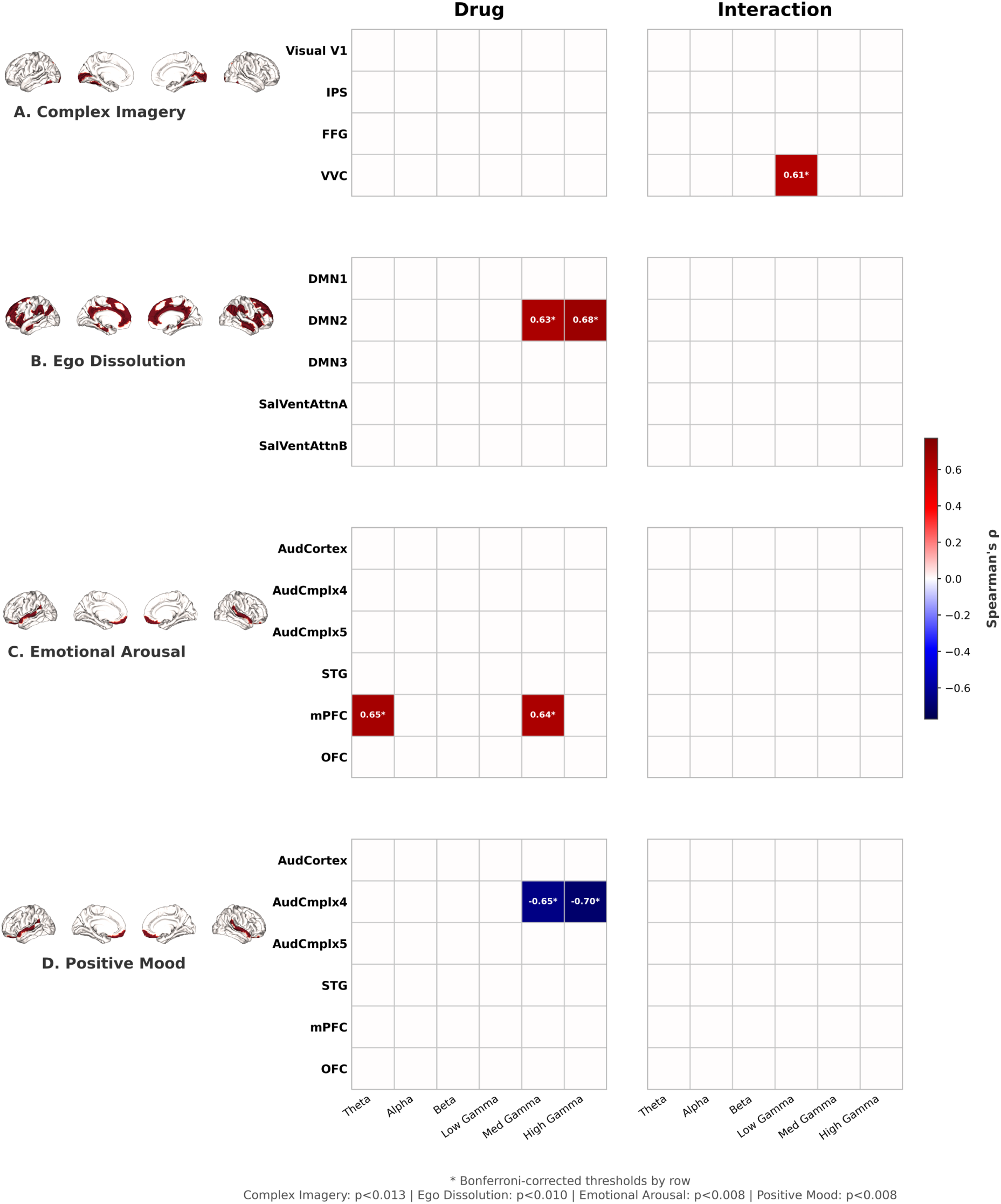
Relation between changes in SDI and Phenomenological dimensions. SDI changes related to **A** Complex Imagery, **B** Ego Dissolution, **C** Emotional Arousal and **D** Positive Mood across frequency bands and statistical conditions (e.g. Main Effect Drug). Spatial maps corresponding to the regions considered were presented above the panel labels. Significance level was set to *p <* 0.05, and multiple-comparison correction was controlled using bonferroni correction per statistical condition within each frequency band. The corrected p-value is displayed underneath the heatmap. Legend: IPS -Intra-parietal Sulcus; FFG -Fusiform Gyrus; VVG -Ventral Visual Gyrus; STG -Superior Temporal Gyrus; mPFC -medial Prefrontal Cortex; OFC -Orbital Frontal Cortex; Yeo Network v17 -Network 7, 8, 15, 16, 17

For Complex Imagery (Fig. 5A), analyses focused on primary visual cortex (V1), the intraparietal sulcus (IPS), the fusiform gyrus (FFG), and ventral visual cortex (VVC). SDI changes within these regions were examined for correspondence with the reported complex imagery across frequency bands and statistical contrasts. Notably, a strong positive correlation (*ρ* = 0.61, bonferroni-corrected; 95% CI : [0.09, 0.86]) was observed for the drug and music interaction in VVC within the low-gamma band, indicating that LSD-induced, music-modulated decoupling in ventral visual regions were associated with enhanced complex visual imagery.

For Ego Dissolution (Figure 5B), analyses targeted large-scale functional networks defined by the 17-network Yeo–Krienen parcellation [31], with a focus on the Default Mode Network (DMN) (three subnetworks) and the salience/ventral attention network (two subnetworks). Stronger decoupling within the DMN, subnetwork 2 — encompassing the Posterior Cingulate Cortex (PCC) and medial prefrontal cortex (mPFC) — showed strong correlations with ego dissolution in the mid- and high-gamma bands under the main effect of drug (*ρ* = 0.63, 95% CI : [0.16, 0.87] and 0.68, 95% CI : [0.26, 0.88], *p <* 0.05, bonferroni-corrected). These associations indicate that LSD-induced decoupling within core DMN regions is robustly associated with the intensity of ego dissolution.

For Emotional arousal and Positive Mood (Figure 5C-D), analyses targeted auditory and prefrontal regions, including primary auditory cortex (AudCortex), auditory complexes 4 and 5 (AudCmplx 4/5), superior temporal gyrus (STG), medial prefrontal cortex (mPFC), and orbitofrontal cortex (OFC). LSD-induced changes in SDI were associated with both emotional arousal and positive mood. Specifically, stronger decoupling in the mPFC was associated with increased emotional arousal across theta (*ρ* = 0.65, 95% CI : [0.14, 0.90], *p <* 0.05, bonferroni-corrected) and mid-gamma bands (*ρ* = 0.64, 95% CI : [0.23, 0.89], *p <* 0.05, bonferroni-corrected). In contrast, increased coupling within AudCmplx 4 correlated with higher positive mood in the mid- (*ρ* = −0.65, 95% CI : [-0.86, -0.25], *p <* 0.05, bonferroni-corrected) and high-gamma (*ρ* = −0.70, 95% CI : [-0.90, -0.32], *p <* 0.05, bonferroni-corrected) ranges.

## 4 Discussion

We utilized connectome harmonics to show that LSD induces robust decoupling of low-frequency activity from anatomical structure, alongside frequency-specific increases in coupling within temporal cortices at higher frequencies. Functional correlates further reveals that these effects are organized in a cognitive- and affective system–specific manner. Finally, inter-individual variability in SFC reorganization relates to subjective experience, including visual imagery, ego dissolution, emotional arousal, and positive mood.

### 4.1 Desynchronization in time is re-expressed across spatial scales

Connectome harmonics [7, 8, 32] enable direct characterization of structure-function organization across spatial scales. By incorporating indirect, polysynaptic pathways, this approach captures dominant patterns of activity propagation beyond direct anatomical connections. As in most SFC studies, however, the topology considered is restricted to cortico-cortical connectivity.

Many studies have utilized structural connectome and its harmonics to investigate SFC primarily in the normal wakeful state [8, 9, 10, 12, 11, 13, 14, 15]. Applications of this framework to psychedelic states have relied largely on fMRI [16, 17, 18], reporting redistribution of BOLD power from spatially smooth to more distributed eigenmodes. While often interpreted as reflecting increased dynamical flexibility, such findings are difficult to disentangle from vascular and metabolic influences [20]. Here we characterize for the first time how neural activity measured with source-localized MEG reorganizes its alignment with the underlying structural connectome under LSD.

Consistent with prior electrophysiological work [33, 34], LSD produced broadband low-frequency attenuation (theta–beta) and increased gamma power, alongside elevated temporal signal diversity [24]. When projected onto connectome harmonics, this temporal desynchronization manifests as a global reduction in graph power across spatial modes (Fig. 2). Because the graph Fourier transform preserves signal energy, reduced oscillatory amplitude necessarily entails reduced total graph power [35]. Thus, gPSD attenuation represents the structural-domain expression of classical low-frequency desynchronization. These effects arise from fast electrophysiological dynamics and should not be directly equated with graph-spectral findings derived from BOLD signals. Exploratory analyses suggested that music attenuated LSD-related graph power changes, although the interaction was not statistical significant, mirroring previously reported music-related modulations of spectral and temporal features in the same dataset [36, 24].

### 4.2 LSDDecouples alpha and beta activity from structural architecture

Given that LSD induces robust low-frequency desynchronization (Fig. S2), a concomitant weakening of graph power (Fig. 2) is expected. The more informative question is whether LSD merely attenuates the contributions of connectome harmonic modes globally, or selectively reweights how neural activity is distributed across the structural connectome. To address this, we used SDI [8], which quantifies the relative contribution of localized (higher-order) harmonics over spatially smooth (lower-order) harmonics.

Under normal wakeful conditions, fMRI-based SFC follows a unimodal–transmodal hierarchy [8], paralleling large-scale functional organization [29]. However, recent electrophysiology-derived SFC revealed modality-specific differences [14, 15] : while unimodal cortices show consistent coupling across modalities, electrophysiology-derived SFC show a coupling-decoupling gradient rather than SFC of BOLD typically occupying the decoupling gradient [14, 11, 15]. These distinctions are reinforced by direct comparisons of MEG- and fMRI-derived SFC [15] and are consistent with evidence that BOLD responses may dissociate from underlying oxygen metabolism [19].

Against this backdrop, examining SFC with direct neural measures provides a less ambiguous characterization and access to a broader frequency range. Within this framework, we find that LSD robustly increases decoupling of low-frequency activity, particularly in the theta–beta range. Low-frequency rhythms reflect large-scale synchronous population dynamics [21, 22] and are thought to support long-range communication [37, 22], rendering them typically well aligned with anatomical pathways [38, 11, 15]. Disruption of large-scale coordination under LSD would therefore be expected to preferentially affect these rhythms. The observed low-frequency decoupling is consistent with this account and suggests selective weakening of structure-aligned integration.

More broadly, theta–beta decoupling reflects altered alignment between functional dynamics and anatomical constraints. Such loosening of structural constraints may contribute to the emergence of atypical brain states characteristic of psychedelic experience. This interpretation aligns with the notion of an expanded repertoire of accessible brain states under psychedelics [16, 2].

### 4.3 Strengthening of bottom-up processes in gamma may underlie psychedelic-induced coupling

Previous work has demonstrated a largely consistent SFC topography across MEG frequency bands during the normal wakeful state [14, 11]. Under LSD, however, SFC reorganizes in a frequency-selective manner. Low-frequency bands (theta–beta) predominantly exhibit decoupling, whereas high-frequency bands (gamma) display a mixture of coupling and decoupling effects (Fig. 3), with coupling most prominently localized to temporal cortices. Across-frequency similarity analyses support this clustering, with low-gamma occupying an intermediate, transitional position (Supp. Fig. S3). This dissociation is broadly consistent with hierarchical models of cortical communication, in which alpha–beta rhythms are associated with top-down predictive control, while gamma activity is associated with bottom-up, stimulus-driven processing via cross-frequency interactions [39]. From this perspective, the REBUS framework [40] offers a useful interpretative context: psychedelics relax high-level priors and top-down constraints, thereby overweighting bottom-up sensory and perceptual processes. Consistent with this account, LSD appears to weaken structure-imposed constraints on slow, integrative coordination mediated by theta–beta rhythms, while preserving—or in selected regions enhancing—structure-aligned coordination in faster gamma dynamics. Although gamma oscillations are often locally generated, evidence suggests they can contribute to long-range communication under specific conditions [41], rendering such coupling effects physiologically plausible.

### 4.4 LSD reorganizes SFC in a cognitive and affective system-specific manner

The functional relevance of LSD-induced SFC changes was assessed using NiMARE. Analyses focused on the main effect of drug (Fig. 4; see Supplementary Fig. S6 for the Drug × Music interaction and main effect of music). Robust spatial effects were observed only for the main drug effect (Fig. 3, cluster-corrected), whereas interaction and music effects were nominal (*p <* 0.05, uncorrected). Because NiMARE operates on unthresholded statistical maps, results for the latter conditions are interpreted cautiously.

Functional correlates of the SDI patterns were assessed in the alpha and high-gamma bands, which capture the two dominant frequency regimes identified in the SFC analyses (Supplementary Fig. S3). Rather than inducing a global shift toward decoupling, LSD produces a selective rebalancing of SFC across cognitive and affective systems, arguing against a unitary account of psychedelics-induced “disorganization” often inferred from BOLD-based studies [16]. Specifically, LSD strengthens SFC within auditory systems while inducing pronounced decoupling within visual systems across both alpha and high-gamma bands. Although this pattern is shared across frequencies, auditory coupling is especially prominent in the high-gamma bands, where effects localize to temporal cortices. In contrast, visual systems consistently occupy the decoupling end of the SDI gradient. This interpretation aligns with prior findings linking LSD-induced visual phenomena to reduced occipital alpha power and increased functional connectivity of primary visual cortex, consistent with disinhibition and expanded influence rather than functional breakdown [23]. In this context, visual decoupling may reflect a relaxation of structural constraints that facilitates internally generated imagery. Supporting this view, mental imagery–related topics are preferentially associated with decoupled regions in the alpha band, consistent with reports linking alpha reductions to both simple and complex imagery under LSD [23].

### 4.5 Changes in structure-function reorganization relates to phenomenological reports

To identify neural correlates of psychedelic experience, we examined whether LSD-induced alterations in SFC relate to phenomenological reports. Rather than adopting a whole-brain exploratory approach, we focused on *a priori* regions and networks consistently implicated in psychedelic and music neuroscience. This targeted strategy reduces multiple-comparisons burden and facilitates interpretation within an established neurobiological framework. However, a wide CI range revealed by bootstrapping analyses warrant a careful interpretation of these findings.

Visual imagery under psychedelics has repeatedly been linked to early and ventral visual cortices. Carhart-Harris and colleagues [23] reported that increased blood flow in primary visual cortex (V1) predicted complex imagery under LSD, while more recent effective-connectivity work under psilocybin implicates a distributed circuit involving early visual cortex, ventral visual regions (including fusiform gyrus), IPS, and inferior frontal gyrus, with imagery related to altered top-down influences [42]. Guided by this literature, we examined whether changes in SFC within these regions tracked imagery ratings. We observed that drug × music–related changes in SDI within VVC were positively associated with complex imagery. Notably, this relationship emerged despite the absence of a statistically significant group-level interaction effect on SDI, indicating that individual differences in music-modulated decoupling scale with imagery vividness even when the interaction is not robust at the group level.

Disruptions in the sense of self under psychedelics have been consistently associated with changes in the DMN and salience network [23, 43]. Prior electrophysiological studies further suggest that reductions in low-frequency oscillatory power (delta and alpha) within DMN regions predict ego dissolution [23]. Extending this literature, we show that SDI derived from electrophysiological signals within core DMN regions robustly predicts ego dissolution in the mid- and high-gamma bands. This finding suggests that a reorganization of how neural activity decouples from the underlying structural scaffold of transmodal networks could potentially manifest a neural correlate predictive of ego dissolution.

Emotional responses to music and psychedelics have been linked to temporal association cortices, mPFC, and ventromedial prefrontal regions [44], with emotional arousal and positive mood showing substantial covariance [45]. Anchoring our analyses in this framework, we related SDI changes in auditory and prefrontal regions to emotional arousal and positive mood. We found that stronger decoupling of mPFC tracked increased emotional arousal across theta and mid-gamma bands, whereas stronger coupling within auditory association cortex (Auditory Complex 4) in mid- and high-gamma bands predicted higher positive mood. These dissociable relationships suggest that emotional intensity and affective valence may rely on distinct structure–function regimes, with emotional arousal linked to reduced structural constraint in prefrontal regions and positive mood linked to strengthened coupling in auditory cortices.

## 5 Limitations and Perspectives

We report the limitations here as it may guide future research. First, this study tackles structure-function coupling of cortical activity that includes both rhythmic (periodic) and arrhythmic (aperiodic) activity. LSD induces widespread alterations in the aperiodic component of the power spectrum [24]. Consequently, the observed SFC changes cannot be attributed exclusively to oscillatory dynamics. This distinction is non-trivial: synchronous rhythmic activity and asynchronous aperiodic activity may exhibit fundamentally different relationships to the anatomical scaffold. Disentangling the structure–function coupling of periodic and aperiodic components – potentially by computing SFC separately on parametrized spectral components – represents an important avenue for future work. Such analyses would clarify whether the observed decoupling primarily reflects disrupted large-scale synchrony, altered excitation–inhibition balance, or a combination of both. Second, the sample size of N = 17, while comparable to other controlled psychedelic neuroimaging studies, limits the precision of effect size estimates. Brain-phenomenology correlations in particular should be interpreted as preliminary, as reflected in the wide bootstrap confidence intervals reported. Replication in larger samples remains necessary to establish generalizability and more precisely characterize the magnitude of LSD-induced structure–function reorganization. Finally, the subjective ratings were collected retrospectively and referenced to participants’ perceived peak drug effects, whereas in-scanner ratings were acquired approximately 150 minutes after the peak. This mismatch likely introduces the noise and reduces the sensitivity of brain-phenomenology correlation. Future studies would benefit from a denser, temporally aligned sampling of phenomenology – ideally collected concurrently with neural recordings – to better capture dynamic brain–experience coupling and improve statistical power.

## 6 Conclusions

By mapping source-localized MEG activity onto connectome harmonics, this study provides evidence for how LSD reshapes structure-function relationships in the human brain. At the level of temporal dynamics, we replicated the canonical signature of the psychedelic state characterized by widespread attenuation in low-frequency bands (theta–beta) alongside increases in gamma power. Re-expressing these temporal signals in the spatial domain using harmonics of the structural connectome revealed a corresponding reduction in graph power across theta–beta and a reversal in mid- and high-gamma. This finding demonstrates how classical desynchronization of temporal dynamics organize in the spatial domain along the anatomical scaffold. Crucially, beyond global power changes, LSD induced a frequency-selective reorganization of structure–function coupling. Low-frequency activity exhibited robust decoupling from the structural connectome, whereas higher frequencies showed a more heterogeneous pattern, including strengthened coupling within temporal cortices alongside focal decoupling elsewhere. Functional correlates of these effects identified using NiMARE revealed a system-level reconfiguration rather than uniform disorganization: LSD preferentially induced decoupling in visual and attention/executive systems while strengthening coupling in auditory and affective/language-related domains. Importantly, these alterations in structure–function organization were meaningfully related to subjective experience. Default-mode decoupling predicted ego dissolution and sensory/prefrontal SDI changes predicted imagery and affective reports. Together, these findings demonstrate that LSD induces decoupling and coupling of structure-function alignment in a system-, frequency-specific manner, and predicts subjective experience.

## 7 EthicsStatement

We utilized the subset of the data collected by Carhart-Harris and colleagues [23]. This original study was approved by the National Research Ethics Service Committee London-West London and was conducted in accordance with the revised declaration of Helsinki (2000), the International Committee on Harmonization Good Clinical Practice guidelines, and the National Health Service Research Governance Framework. Imperial College, London sponsored the research, which was conducted under a Home Office license for research with Schedule 1 drugs.

## 8 Data and Code Availability

Queries regarding data are to be directed to SM and RC-H. Code is available on https://github.com/venkateshness/LSD_SDI

## 9 AuthorsContributions

**VS** : Conceptualization, Methodology, Software, Validation, Formal Analysis, Investigation, Data Curation, Writing - Original Draft, Writing - Review & Editing, Visualization. **AP** : Conceptualization (Preprocessing, Source Localization), Methodology (Preprocessing, Source Localization), Writing - Review & Editing. **JB** : Conceptualization (Statistics), Methodology (Statistics), Writing - Review & Editing. **YH** : Conceptualization (Preprocessing), Writing - Review & Editing. **SM** : Data Curation, Writing - Review & Editing. **RC-H** : Data Curation, Writing - Review & Editing. **KJ** : Conceptualization, Writing—Review & Editing, Supervision, Project Administration, Funding Acquisition; **GL** : Conceptualization, Methodology, Resources, Writing—Review & Editing, Supervision, Project Administration. **NF** : Conceptualization, Methodology, Resources, Writing—Review & Editing, Supervision, Project Administration, Funding acquisition.

## 10 Declarations of Competing Interests

RC-H reports providing scientific advice for TRYP therapeutics, Osmind, Otsuka, and Red Light Holland. Others declare no competing interests.

## Acknowledgments

We thank several colleagues who contributed to the progress of this study. We are grateful to Jordan O’Byrne for assistance with dataset organization; Kenneth Shinozuka for identifying missing data for two subjects; and Yassine El Ouahidi for adapting the functional decoding pipeline from Neurosynth to NiMARE. Our Qsirecon pipeline was run in a High-Performance Cluster grid thanks to the BC DRI Group, Calcul québec, and the Digital Research Alliance of Canada https://alliancecan.ca/, and we really appreciate their continued support.

K.J. is supported by funding from the Canada Research Chairs (950-232368) program and a Discovery Grant from the Natural Sciences and Engineering Research Council of Canada (2021-03426).

We utilized a subset of the data collected by Carhart-Harris and colleagues [23]. Data collection of this study was supported by Safra Foundation and the Beckley Foundation as part of the Beckley-Imperial research programme, and by supporters of the Walacea.com crowdfunding campaign.

## 11 Declarationof LLM Usage

We used OpenAI’s ChatGPT 5.2 to improve writing style, fluidity, and brevity. All AI-suggested content was reviewed and integrated into the manuscript when appropriate. The authors take full responsibility for the manuscript content.

## 12 Supplementary Material

Below contain details about the methodology of preprocessing MEG signal, and Structural Connectome reconstruction. Furthermore, a full mathematical description to quantify structure-function coupling is provided. Supporting results guide our interpretation, alongside additional exploratory analysis of main effect music and interaction effect drug x music.

### 12.1 MEGPreprocessing

MEG preprocessing followed the same general pipeline as in the original dataset [23], with the addition of automated artefact detection procedures implemented using the Autoreject package [25]. A schematic overview of the preprocessing workflow is provided in Fig. S1. In brief, continuous recordings were filtered, visually inspected, and segmented into non-overlapping 2-s epochs. Artefacts were then attenuated through a two-stage procedure combining automated epoch rejection and independent component analysis (ICA). Specifically, an initial pass of Autoreject was used to identify and mask artifactual epochs using data-driven, sensor-specific thresholds prior to ICA decomposition. ICA (Picard algorithm [46]) was then applied to remove components associated with physiological noise, after which cleaned signals were reconstructed by back-projecting the retained components. A second pass of Autoreject was subsequently applied to the reconstructed data to further attenuate residual artefacts, yielding the final cleaned epochs used for all subsequent analyses. Following this pipeline, approximately 200 clean epochs per recording on average (SD = 20) were available for subsequent analyses, with comparable data retention across LSD and PLA conditions (*t* = −1.69, *p <* 0.12).

### 12.2 SourceLocalization

High-resolution T1-weighted anatomical MRI scans were available for all participants and sessions [23]. Structural images were processed using FreeSurfer’s standard reconstruction pipeline (recon-all, v7.4.1), which includes motion correction, intensity normalization, skull stripping, and white–gray matter surface segmentation. All reconstructions were visually inspected for accuracy prior to source analysis. Subject-specific cortical surfaces were used to constrain MEG source estimation. Forward models were computed using Boundary Element Model (BEM) surfaces generated with FreeSurfer’s watershed algorithm as implemented in MNE-Python [47]. A standard single-layer (inner-skull) BEM was used with conductivity parameter set to 0.3. Individual source spaces were defined using an ico5 tessellation, yielding 10242 vertices per hemisphere. MEG sensor positions were co-registered to individual anatomical images using three fiducial points (nasion, left and right preauricular points), resulting in an affine transformation mapping MEG sensor coordinates to MRI space. Inverse solutions were computed using dynamic Statistical Parametric Mapping (dSPM) [48]. Because empty-room noise recordings were unavailable, noise covariance was modeled using an identity matrix, assuming uniform sensor noise. Source orientation constraints were set to be mostly perpendicular to the cortical surface, adopting a loose orientation model (loose = 0.2). Depth weighting was applied to the forward model with a depth prior of 0.8. This procedure yielded source-level time series at each vertex of the cortex.

For group-level analyses, individual source estimates were morphed to the FreeSurfer fsaverage (v5) template. Morphed source time series were subsequently parcellated onto the HCP-MMP atlas [26] using MNE’s extract_label_time_course function. Regional activity was obtained by averaging across vertices within each parcel, resulting in 360 cortical region time series per participant and condition. Power Spectral Density (PSD) estimates were computed at the source level using multitaper method (implemented in MNE’s compute_psd_epochs), followed by performing source space morphing to extract the PSDs in HCP-MMP ROIs.

Source inversion was performed on epochs basis (Clean Signal; Fig. S1). Because these epochs are 2-s windows, performing bandpass filtering risks edge artefacts. We therefore padded zeros of duration 1 second on both side creating 4-s epochs to apply filters (butterworth, order of 4): theta (4 - 8Hz), alpha (8 - 13 Hz), beta (15 -30 Hz), low-gamma (30 -60 HZ), mid-gamma (60 -90 Hz) and high-gamma (90 -120 Hz).

### 12.3 StructuralConnectome Reconstruction and Structure-Function Coupling

We derived the structural connectome from high-quality diffusion MRI data provided by the Human Connectome Project (HCP), since diffusion-weighted imaging (DWI) was not available for our dataset. We used the S1200 data release, comprising 1,063 healthy participants with preprocessed diffusion and T1-weighted anatomical images [27]. Diffusion data were processed using a state-of-the-art tractography pipeline implemented in QSIRecon [49] (automated report below):

#### Automated Report from NiReports (Excerpt)

Reconstruction was performed using *QSIRecon* 1.1.1.dev0+gaf43da9.d20250414 [49], which is based on *Nipype* 1.9.1 ([50]; [51]; RRID:SCR_002502).

A hybrid surface/volume segmentation was created [52]. FreeSurfer outputs were registered to the QSIRecon outputs.

##### Anatomical data for DWI reconstruction

T1w-based spatial normalization calculated during preprocessing was used to map atlases from template space into alignment with DWIs. Brain masks from antsBrainExtraction were used in all subsequent reconstruction steps. The following atlases were used in the workflow: the glasser atlas. Cortical parcellations were mapped from template space to DWIs using the T1w-based spatial normalization.

##### MRtrix3 Reconstruction

Multi-tissue fiber response functions were estimated using the dhollander algorithm. FODs were estimated via constrained spherical deconvolution [53, 54] using an unsupervised multi-tissue method [55, 56]. Reconstruction was done using MRtrix3 [57]. FODs were intensity-normalized using mtnormalize [58].

Many internal operations of *QSIRecon* use *Nilearn* 0.10.1 [59] and *Dipy* 1.8.0 [60]. For more details of the pipeline, see [the section corresponding to workflows in *QSIRecon*’s documentation](https://qsirecon.readthedocs.io/en/latest/workflows.html).

In line with previous SFC studies [8, 14], we defined the adjacency matrix of the structural connectome as the density of fibers connecting pairs of HCP-MMP ROIs, same atlas in which sources of the MEG activity are estimated.

#### 12.3.1 ConnectomeHarmonic Decomposition and Structural Decoupling Index

The harmonics of the structural connectome are derived using Eigendecomposition [7]. Let us define an undirected weighted graph *G* =*< V,* E *, W >*, where *V* is a set of *N* elements called vertices (i.e., 360 HCP-MMP ROIs) and E ⊂ *V* × *V* the set of edges connecting unordered pairs of vertices with scalar weights *w* (i.e., fiber density). The degree matrix **D** ∈ *R^N^*^×^*^N^* of a graph *G*, is a diagonal matrix such that 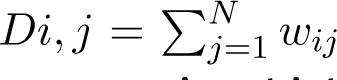 The Adjacency Matrix **A** ∈ *R^N^*^×^*^N^* is the square matrix of dimension *N* × *N* in which each element is different from zero only if the corresponding edge exist. Following previous studies [8, 14], the graph Laplacian is defined as **L** = **I** -**D**^−^**^1^**^/^**^2^ AD**^−^**^1^**^/^**^2^**, with **I** the identity matrix of order *N* . Laplacian matrix **L** is a real symmetric matrix and can be diagonalized as

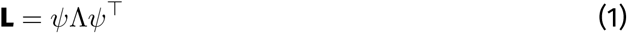

where *ψ* represents the matrix of eigenvectors and Λ the diagonal matrix of real eigenvalues, sorted in increasing order. The eigenvectors are orthogonal to each other and form a complete basis set, which can be leveraged to re-express any spatiotemporal activity [7, 61]. The eigenvectors of the Laplacian of the structural connectome represent the brain’s intrinsic eigenbasis – the fundamental spatial patterns of the connectome. These modes are orthogonal to each other and form a complete basis set, which can be leveraged to re-express any spatiotemporal activity [7, 61]. A subset of these eigenmodes is displayed in Fig. 1B.

Functional activity was mapped onto the structural connectome using Graph Fourier Transform (GFT; [35]), which decomposes the MEG activity *S_t_* into connectome-derived spatial harmonics:

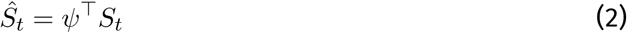

The resulting coefficients *S*^^^*_t_* represent the weights of the contributions of the eigenmodes, describing how neural activity is distributed across connectome harmonics. Squaring these coefficients defines the graph power spectral density (gPSD). The original MEG activity can be reconstructed via the inverse GFT as

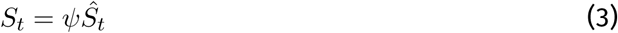

Using the GFT framework, spatial graph filters were defined to decompose the MEG activity into low graph-frequency components (*S*^coupled^), corresponding to spatially smooth activity patterns aligned with the structural connectome, and high graph-frequency components (*S*^decoupled^), corresponding to more spatially localized activity patterns that deviate from large-scale anatomical constraints (see [14] for a detailed formulation).

Following [8], these components were separated using a median split of the gPSD, yielding a critical graph frequency *C* that defines the boundary between low- and high-frequency harmonics. This critical frequency was computed separately for each subject and each temporal frequency band and varied across individuals and spectral bands from theta to gamma (see Supplementary Fig. S4).

The Structural-Decoupling Index (SDI [8]) quantifies the ratio of norm (*l1*-norm) of decoupled activity over coupled activity, and is defined as

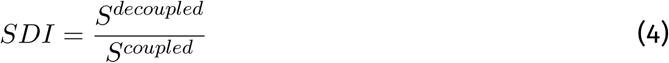

Binary log is applied subsequently, resulting in the positive tail corresponding to decoupling and the negative tail coupling.

### 12.4 FunctionalDecoding of the SDI maps

We performed functional decoding of the SDI contrast maps using NiMARE [28], following established approaches in prior work [29, 8, 14]. The goal of this analysis is to relate the LSD-induced SFC changes to broad cognitive and affective domains, thereby providing a systems-level interpretation of structure–function reorganization under LSD. Functional decoding was performed using the Neurosynth database, a large-scale automated meta-analytic resource comprising activation maps derived from more than 14,000 neuroimaging studies. Neurosynth organizes the neuroimaging literature into topics [62], which correspond to latent semantic components extracted from the co-occurrence of terms in article abstracts and their associated activation patterns. Each topic thus represents a broad cognitive or affective construct (e.g., perception, memory, emotion, language), rather than a specific task or anatomical region. For each SDI contrast (LSD vs. PLA), unthresholded regional SDI maps were rank-ordered and segmented into ten equally sized deciles (10% increments). Each decile was binarized and submitted to the ROIAssociationDecoder implemented in NiMARE, which quantifies the spatial correspondence between the input SDI masks and Neurosynth topic maps. The resulting correlation coefficients reflect the degree to which LSD-induced SDI changes preferentially overlap with brain regions associated with each cognitive or affective topic. Correlation values were converted to z-statistics, and only associations surviving a threshold of *p <* 0.001 were retained. Following previous work [29], topic terms were ranked based on the weighted average of their z-scores across deciles, highlighting the cognitive and affective domains most strongly associated with regions exhibiting the largest LSD-induced alterations in SFC.

### 12.5 LSDinduces desynchronization in the low-frequency bands

LSD-induced modulations in temporal dynamics were revealed by the Fourier analysis across canonical frequency bands: theta (4 -8 Hz), alpha (8 -13 Hz), beta (13 -30 Hz), low-gamma (30 -60 Hz), mid-gamma (60 -90 Hz) and high-gamma (90 -120 Hz). Supplementary Figure S2 shows statistically significant LSD–PLA contrasts (*p <* 0.05, permutation-corrected; 50,000 permutations), with blue indicating power reductions and red indicating power increases.

LSD produced robust reductions in low-frequency power spanning theta through beta. Although this attenuation follows a consistent desynchronization, its spatial extent exhibits clear frequency-specific organization. Theta band power reductions were the most widespread, encompassing large portions of posterior cortex and core regions of default mode network (DMN). Alpha band reductions followed a similar but more spatially restricted pattern, remaining prominent in visual cortex, posterior cingulate cortex (PCC), and temporal regions. Beta band power reductions were more focal, primarily involving anterior and posterior cingulate cortices.

In contrast, higher frequencies exhibited a reversal of this pattern. LSD was associated with significant increases in gamma band power, with low-gamma effects remaining focal, whereas mid- and high-gamma bands showed widespread power increases. The spatial topographies of mid- and high-gamma effects were highly similar, with high-gamma increases extending more prominently into frontal regions (both laterally and medially) and posterior cortical areas.

**13.6 LSD-induced structure–function coupling changes show increasing similarity with temporal frequency proximity**

**13.7 Graph power shifts across temporal frequencies**

**13.8 Nominally significant effects of music interacting with LSD**

**Figure S1:**
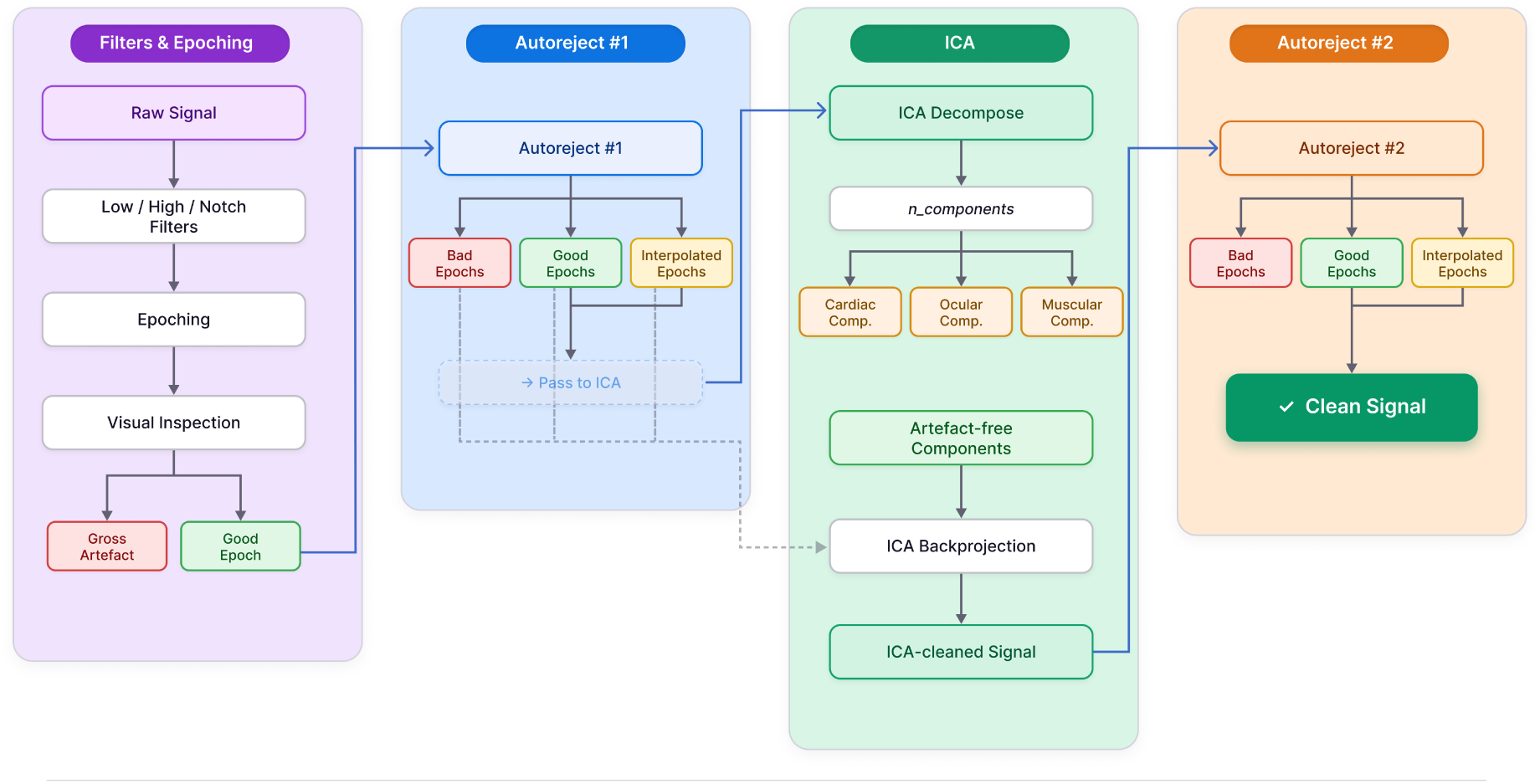
Overview of MEG preprocessing. The pipeline begins with filtering (Bandpass: 1 -120Hz, Notch filter: 50 and 100 Hz), segmenting continuous signal into epochs of 2-s, visual inspection to exclude gross artefacts. The first pass of Autoreject labels the bad epochs, which are then excluded prior to ICA. Artefactual components are excluded through ICA decomposition, followed by backprojecting the artefacts-free components on the output of first pass of Autoreject including bad epochs (purple dashed line). Finally, a second pass of Autoreject is employed to eliminate the residual artefacts. The resulting good and interpolated epochs constitute the *Clean Signal*.

**Figure S2:**
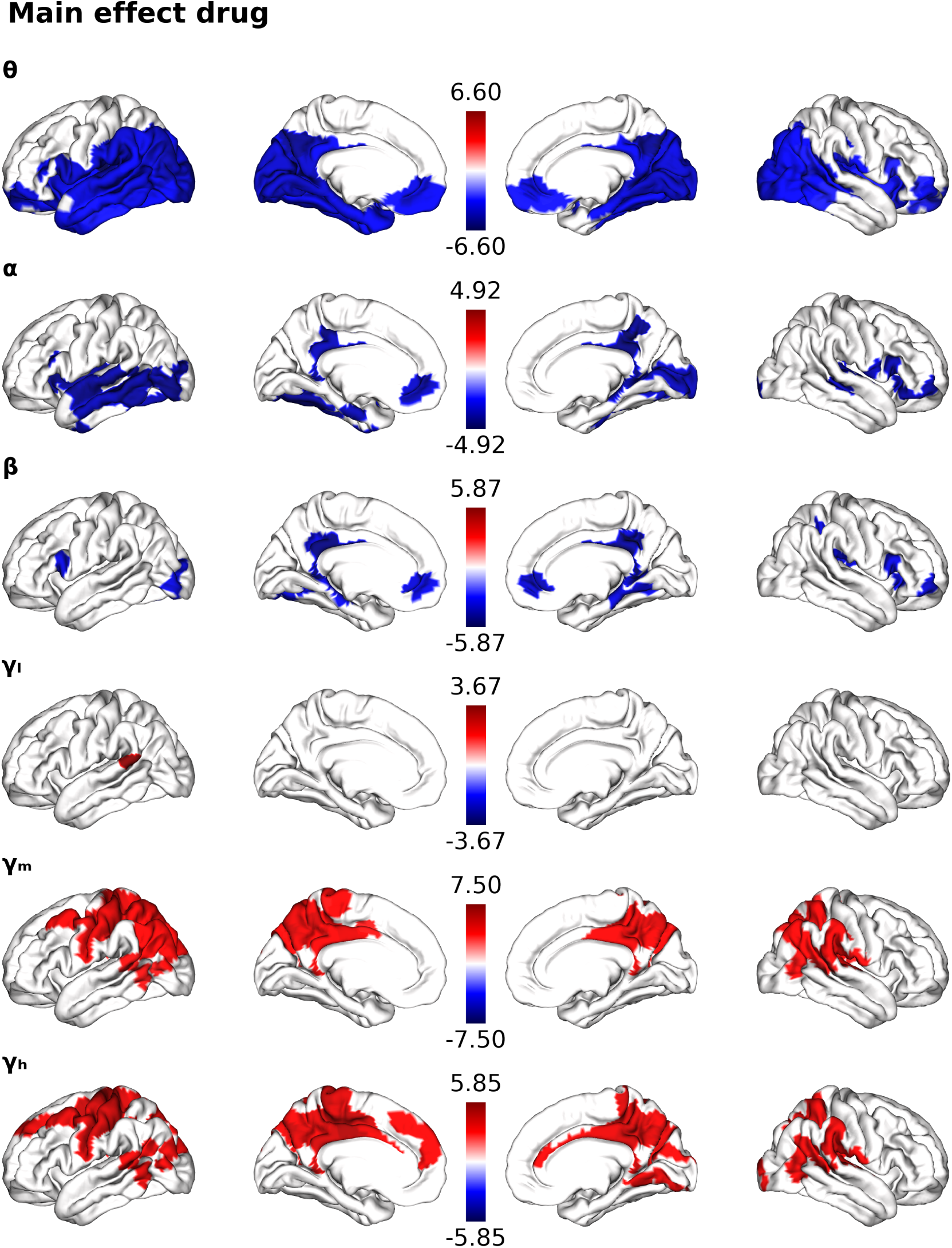
Region-resolved spectral effects of LSD. Significant changes (*p <* 0.05, permutation-corrected, N=50,000) in spectral power (LSD -PLA) across canonical frequency ranges : *θ* (4 -8 Hz), *α* (8 -13 Hz), *β* (15 -30 Hz), low-*γ* (30 -60 Hz), mid-*γ* (60 -90 Hz) and high-*γ* (90 -120 Hz). Blue tail corresponds to LSD-induced weakening of power

**Figure S3:**
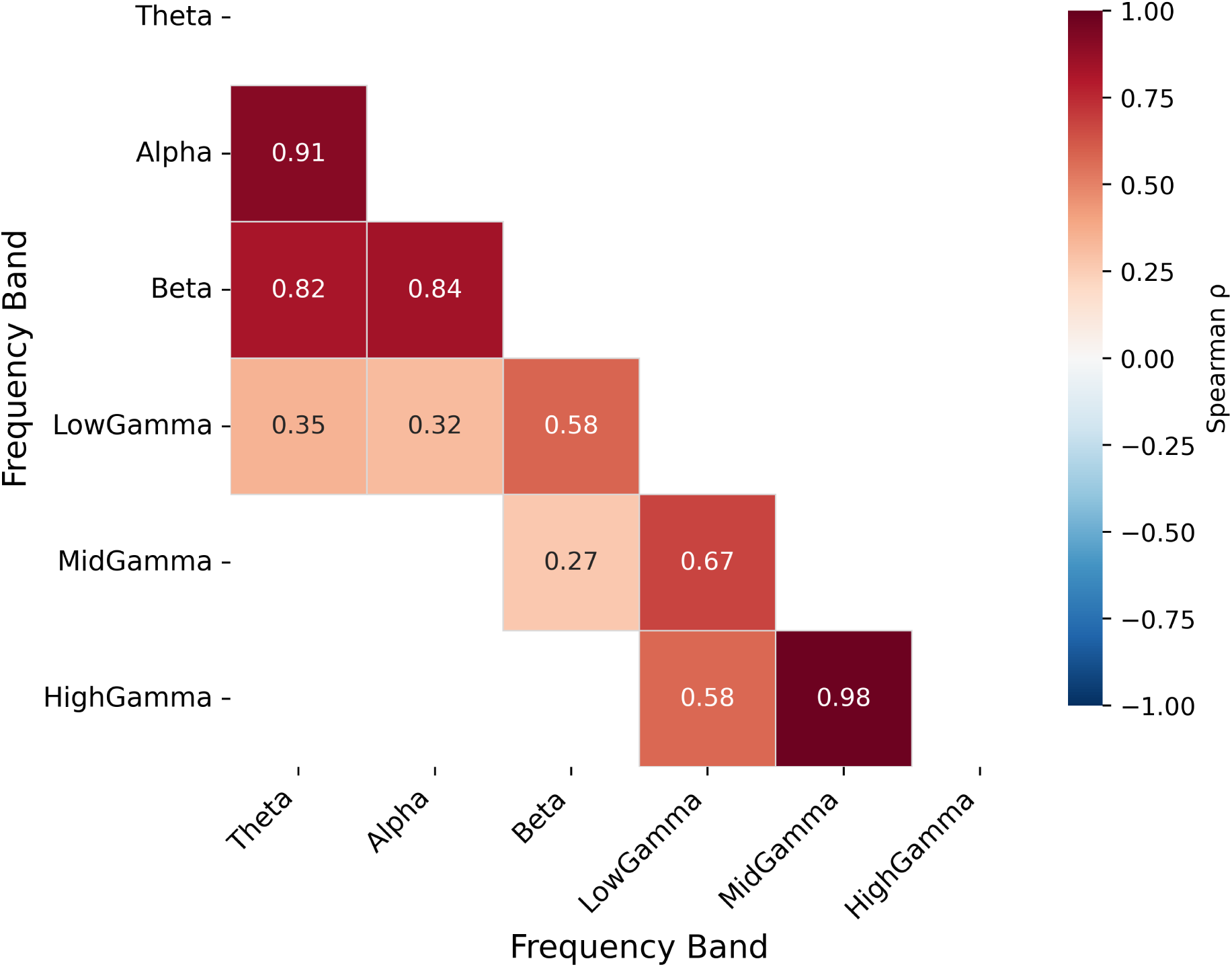
Spatial similarity of SDI Main Effect Drug across frequency Bands. Significant spatial similarity (*p <* 1*e* − 15)) quantified across all pairs by computing the Spearman’s *ρ* on the test statistics resulting from the LSD-PLA contrasts.

**13.9 Nominally significant effects of music**

**13.10 Functional decoding of music and interaction effect maps**

**13.11 Phenomenological scores**

**Figure S4:**
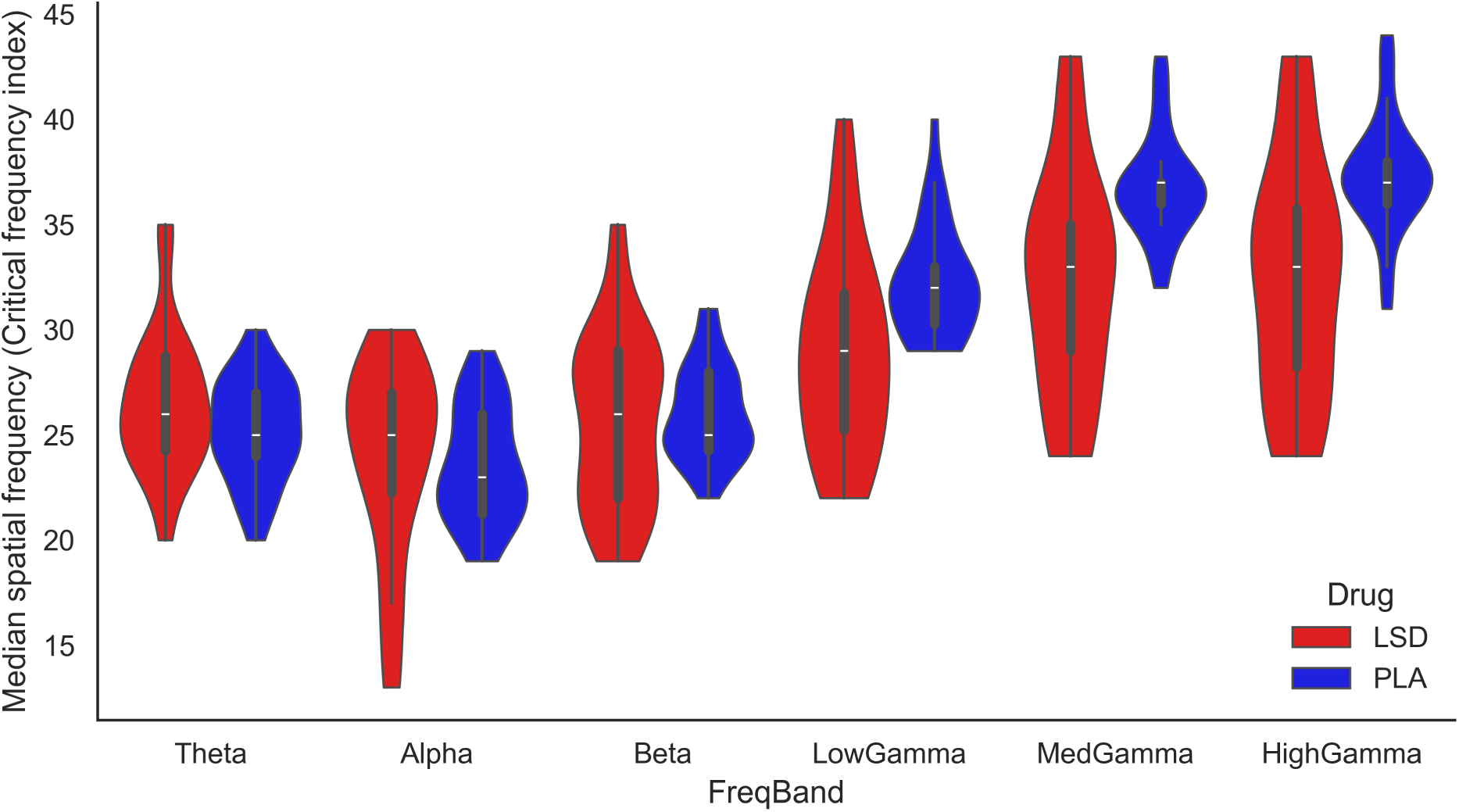
The critical frequency index *C* that splits the graph power spectrum into two equal parts. *C* was used to isolate coupled and decoupled component of the signal.

**Figure S5:**
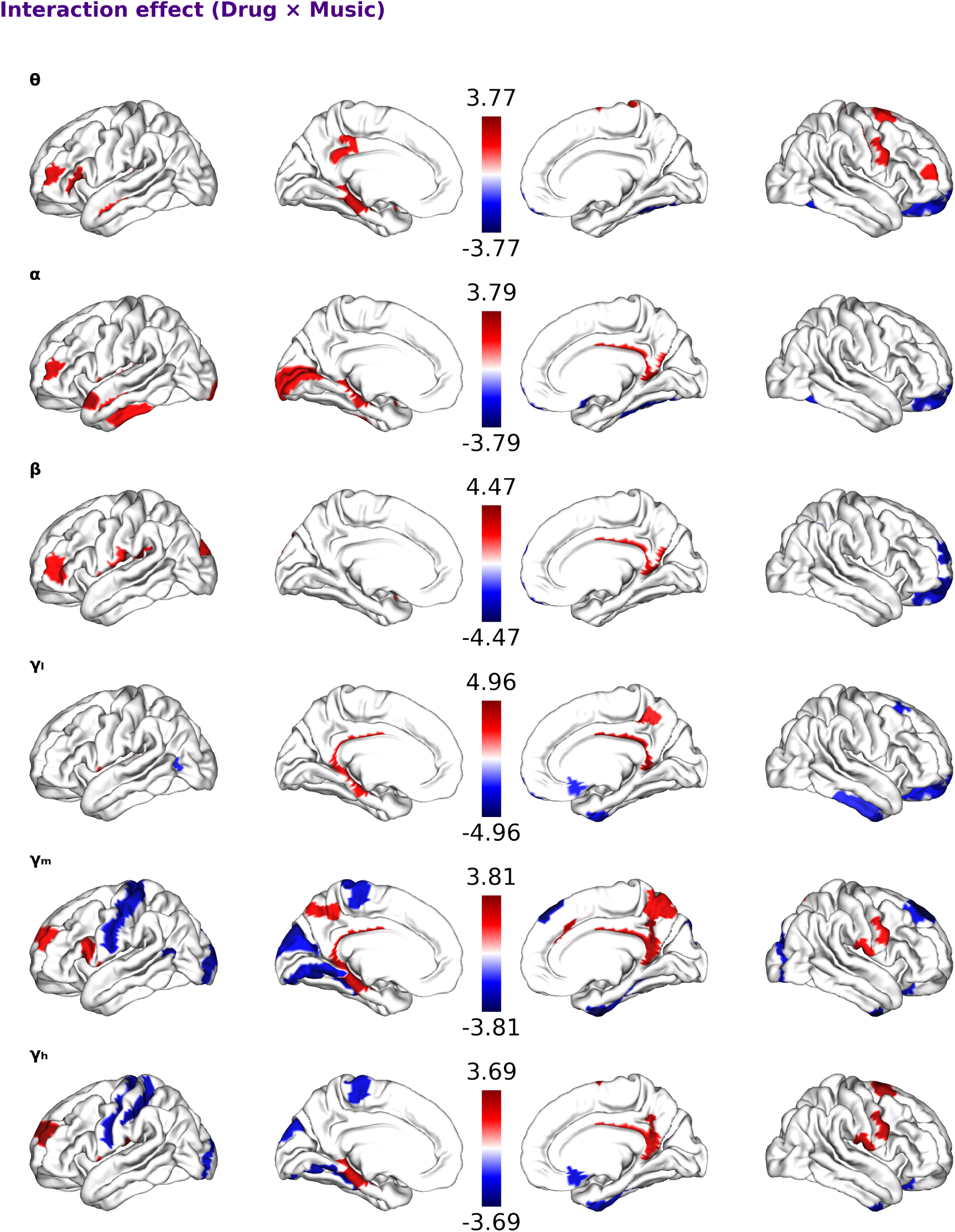
SDI contrast maps (*p <* 0.05, uncorrected) for Interaction Effect.

**Figure S6:**
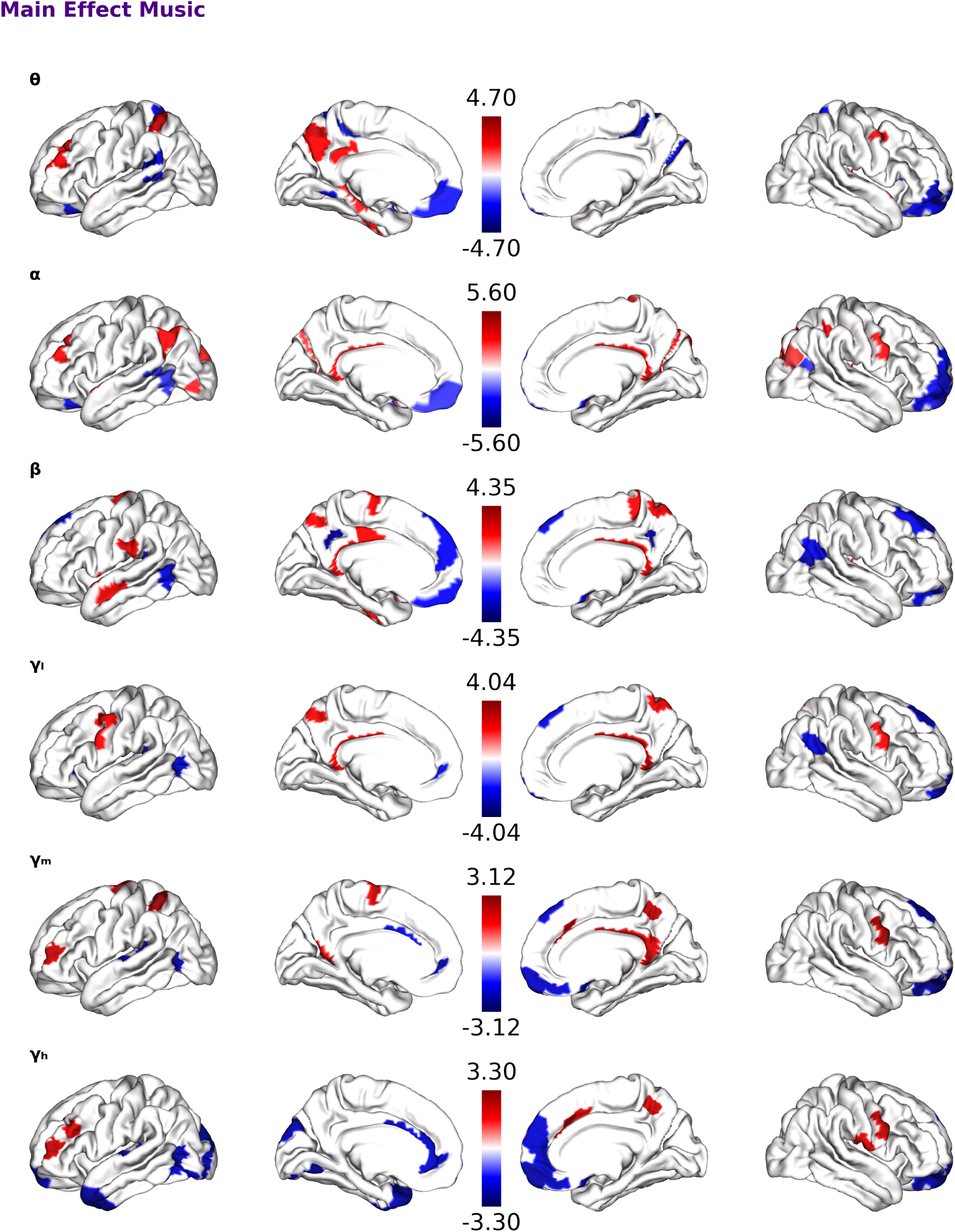
SDI contrast maps (*p <* 0.05, uncorrected) for the effect of music.

**Figure S7:**
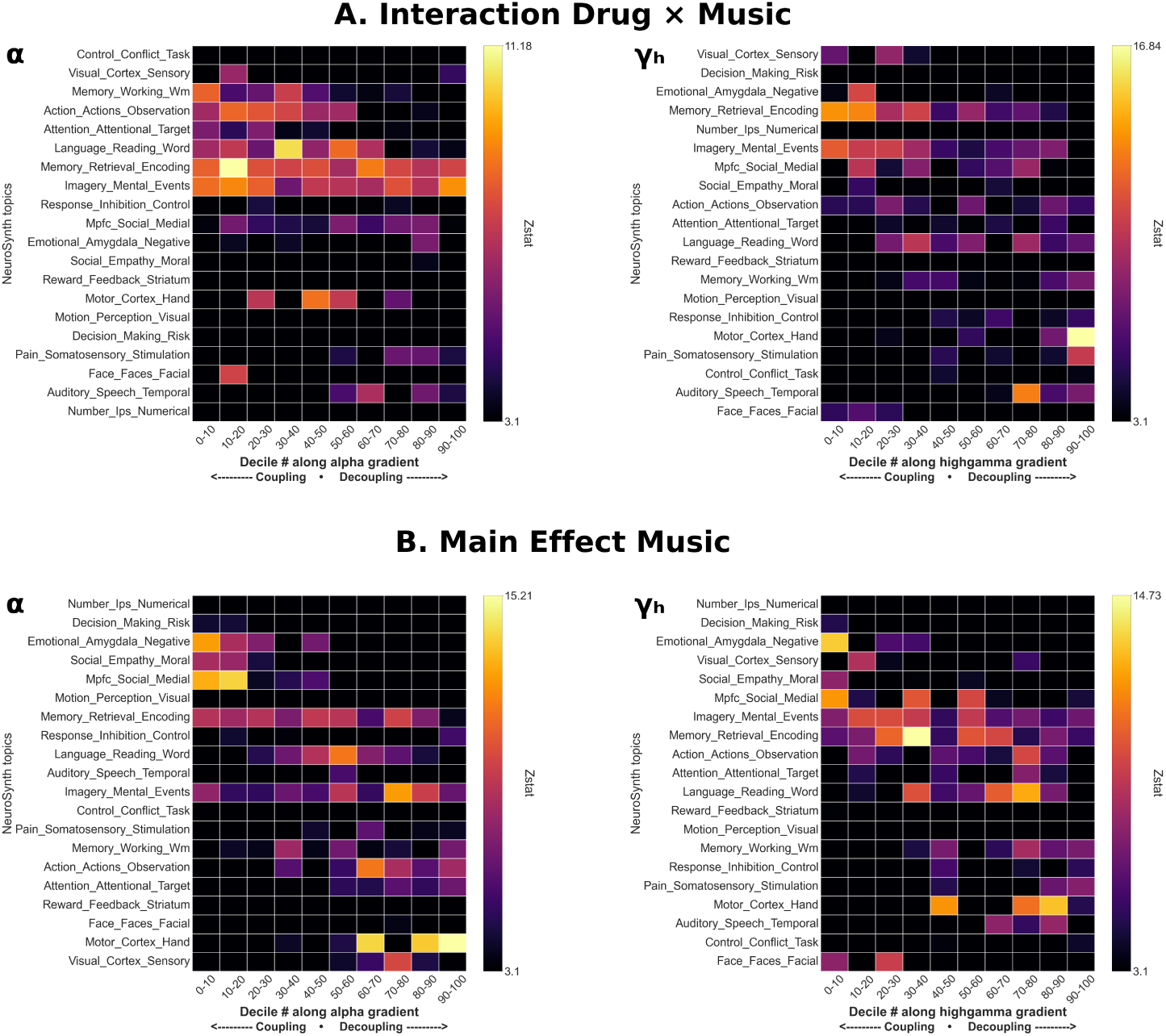
Functional decoding of contrast maps from Main Effect Music and Interaction Effect LSD x music

**Figure S8:**
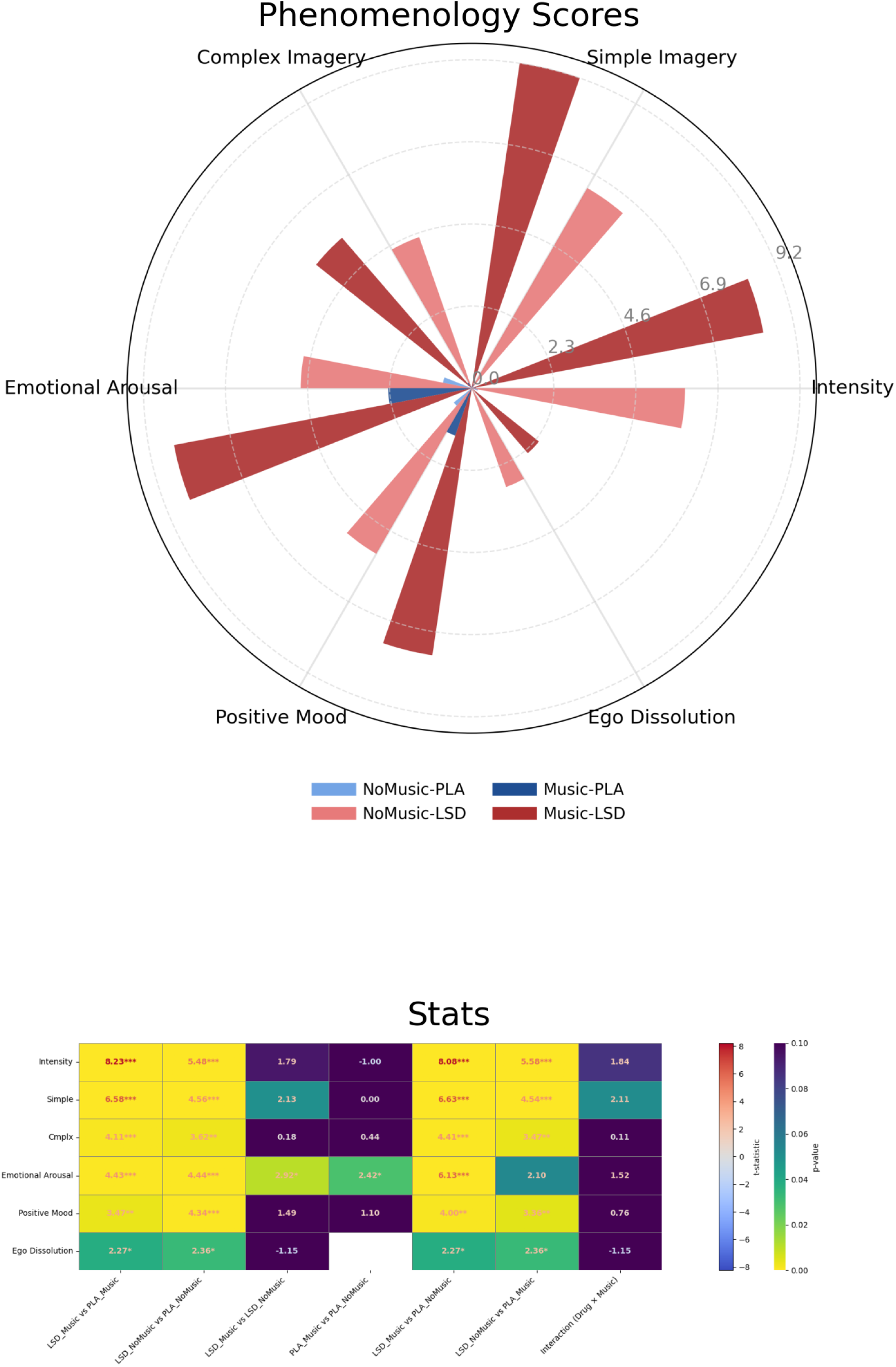
Raw phenomenological scores (top). Statistical comparisons (bottom)

